# Evaluating low-cost hyperspectral and RGB fusion with PSNet for presymptomatic detection of *Albugo candida* in *Arabidopsis thaliana*

**DOI:** 10.64898/2026.03.02.709086

**Authors:** Gabriel U. Crabb, Yiding Zhao, Nicholas K. Priest, Xi Chen, Volkan Cevik

**Author notes:** Corresponding authors: Volkan Cevik; Xi Chen; Nicholas K. Priest. These authors contributed equally to this work (Gabriel U. Crabb, Yiding Zhao).

## Abstract

Plant pathogens cause substantial yield losses worldwide, threatening food security and livelihoods. Because early infection is difficult to diagnose, management often relies on prophylactic pesticide use, increasing costs and environmental impacts. Here we evaluate PSNet, a multimodal framework that fuses hyperspectral and RGB imaging for presymptomatic plant disease detection, together with a low-cost Raspberry Pi hyperspectral camera. We tested the system using *Arabidopsis thaliana* inoculated with the oomycete *Albugo candida* in a controlled experiment comprising 96 plants imaged at 2, 4 and 6 days post inoculation (dpi). PSNet achieved 90.00% overall accuracy on the four-class task comprising Healthy, 2 dpi, 4 dpi and 6 dpi. Collapsing the same true and predicted labels into healthy versus inoculated yielded 97.50% binary accuracy across all dates. Supplementary date-specific binary analyses compared inoculated and mock-inoculated plants separately, using plant-disjoint five-fold cross-validation, three training seeds and within-plant probability aggregation. PSNet achieved balanced accuracies of 55.79%, 83.56% and 95.37% at 2, 4 and 6 dpi, respectively. The 4 dpi observations preceded visible symptoms, whereas at 6 dpi abaxial sporulation was present despite the imaged adaxial surface remaining symptom-free. RGB-only models also performed strongly at 4 and 6 dpi, while HSI-only discrimination weakened after 2 dpi, and no consistent advantage of hyperspectral fusion over RGB alone was demonstrated. Whole-cube spectral analysis identified significant treatment-group differences across 269 of 313 spectral positions at 2 dpi, but none at 4 or 6 dpi after within-time-point Benjamini–Hochberg correction. Because treatment was confounded with tray and acquisition order, these spectral differences cannot be attributed specifically to infection. Together, these results support the use of low-cost multimodal imaging for controlled-condition plant disease classification, while highlighting the need for independent validation before diagnostic deployment.

## 1. Introduction

Plant diseases pose a major and persistent threat to global food security, with pests and pathogens together responsible for up to 40% of annual crop losses worldwide [1, 2]. Climate change and expanding global trade are altering host–pathogen dynamics and increasing the risk of disease emergence and spread [3, 4]. Infection is often recognised only after visible symptoms develop, when intervention may be less effective and yield loss may already have occurred. Earlier detection could enable more targeted management, reduce unnecessary chemical inputs and improve the efficiency of crop protection.

In practice, disease control often relies on prophylactic or calendar-based applications of fungicides and pesticides. Although these approaches can reduce outbreak risk, they increase production costs, contribute to environmental contamination and non-target effects, and accelerate the evolution of chemical resistance in pathogen populations [5, 6]. Reliable presymptomatic diagnostics could therefore support more targeted intervention and reduce blanket chemical use.

Existing diagnostic approaches trade sensitivity against deployability. Visual scouting is subjective and detects disease only after symptoms appear [7]. Molecular assays, including PCR-based methods and isothermal amplification, offer high sensitivity but require specialised equipment, controlled laboratory conditions and technical expertise [8, 9], while CRISPR-based diagnostics remain at an early stage of field validation [10]. Hyperspectral imaging (HSI) offers a non-destructive alternative by recording spectral information across contiguous wavelength bands, enabling detection of changes in pigment content, water status and tissue structure associated with biotic stress [11–14]. Presymptomatic disease detection using HSI has been demonstrated in several host–pathogen systems, including tobacco and tomato [15, 16]. However, wider adoption remains constrained by the cost of commercial systems, the computational demands of hyperspectral data, and variation in performance associated with sensor design, calibration and analytical pipelines [17].

Experimental design is also critical when evaluating plant disease classification. Hyperspectral datasets often contain multiple observations from the same biological individual, so partitioning should be performed at the plant level to prevent closely related samples from appearing in both training and test sets [18]. In longitudinal experiments, comparisons between infection stages can also be confounded by time-related differences in plant age or imaging conditions. Evaluating inoculated and mock-inoculated plants separately within each imaging time point therefore provides a clearer assessment of treatment-group discrimination.

Most hyperspectral classification approaches, from support vector machines [19] to 3D–2D convolutional networks [20], use the hyperspectral cube as their sole input. Combining HSI with inexpensive RGB imaging provides an opportunity to integrate spectral information with higher-resolution spatial and morphological context. Multimodal fusion is therefore a natural extension of hyperspectral classification, but whether the hyperspectral stream contributes information beyond RGB alone during early infection remains less well established.

Here we describe and evaluate a low-cost imaging unit adapted from an open-source 3D-printed hyperspectral camera [21] and used together with RGB imaging in the *Arabidopsis thaliana–Albugo candida* pathosystem. *A. thaliana* is a genetically tractable host with a well-characterised immune system [22], while *A. candida* causes white rust in *Arabidopsis* and *Brassica* crops [23]. Plants were imaged at 2 and 4 days post inoculation (dpi), before visible symptoms, and at 6 dpi, when sporulation was present on the abaxial leaf surface while the imaged adaxial surface remained symptom-free. We aimed to (i) document the imaging unit, its cost and acquisition requirements; (ii) evaluate four-class classification and derive healthy-versus-inoculated accuracy by collapsing the predictions; (iii) compare RGB-only, HSI-only and multimodal models to assess whether the hyperspectral stream contributes additional information; and (iv) define the conclusions that can be supported by this single-run dataset. PSNet, a dual-stream HSI–RGB Transformer [24], was evaluated alongside RGB-only, HSI-only and alternative multimodal baselines, together with ablation variants.

## 2. Materials and methods

Sections 2.1–2.4 describe the biological material, the imaging unit, the acquisition protocol, and the calibration and preprocessing. Section 2.5 defines the primary four-class analysis, the derived binary result and the supplementary date-specific binary evaluation design. Section 2.6 describes PSNet and the implementation used for the supplementary binary comparisons. Section 2.7 describes the supplementary spectral statistics.

### 2.1. Plant material, inoculation and infection validation

The host was the fully susceptible *A. thaliana* line Col-*eds1.2 wrr7* [25], which is highly susceptible to the oomycete *A. candida* (white blister rust) race 4 isolate AcEM2. Seeds were obtained from a certified stock centre. Five 6 × 4 cell trays were filled with a well-drained soil:sand substrate (F2s), and an additional pot with the same mix was used for germination. Seeds were broadcast, vernalised at 4 °C in darkness for 72 h, and transplanted at one plant per cell. Trays stood in water-filled catch trays to maintain uniform moisture by capillarity. Humidity lids were removed at the first true-leaf stage. Unless stated otherwise, conditions were 22 °C, 16 h light/8 h dark, ∼60% relative humidity and 120–150 μmol m^−2^ s^−1^ photosynthetic photon flux density. Seedlings were inoculated about three weeks after transplanting, when rosettes bore four to five true leaves.

Spray guns, flasks and bottles were wiped with 100% ethanol before use. Leaves bearing visible *A. candida* pustules were suspended in chilled sterile water, and sporangia were dislodged by gentle agitation, filtered through nylon mesh and diluted on ice to 10⁵ spores ml⁻¹. Controls received sterile water only. A single run used 96 seedlings (72 inoculated, 24 mock-inoculated). Both leaf surfaces were misted to run-off. Trays were then held at 4 °C for 24 h and placed in a cabinet at 22 °C with 10 h light and 18 °C with 14 h dark until 2 dpi, after which standard growth conditions resumed. Symptoms were assessed visually on the abaxial surface at each imaging time point. To validate latent infection, trypan blue staining was performed at 2, 4 and 6 dpi on a reserved subset of leaves at each time point and examined microscopically as described previously [26]. Staining is described qualitatively; no treatment-specific infection prevalence is inferred from these observations.

### 2.2. Low-cost imaging unit

The hyperspectral camera was adapted from the open-source 3D-printed design of Salazar-Vazquez and Mendez-Vazquez [21]. The build uses a Raspberry Pi Camera Module 3 NoIR, a Raspberry Pi 4B and a 35 mm VIS–NIR lens within the existing optical design (Fig. 2). Detailed modifications and component costs are documented in Appendix S1 and Table S8.

**Figure 1.**
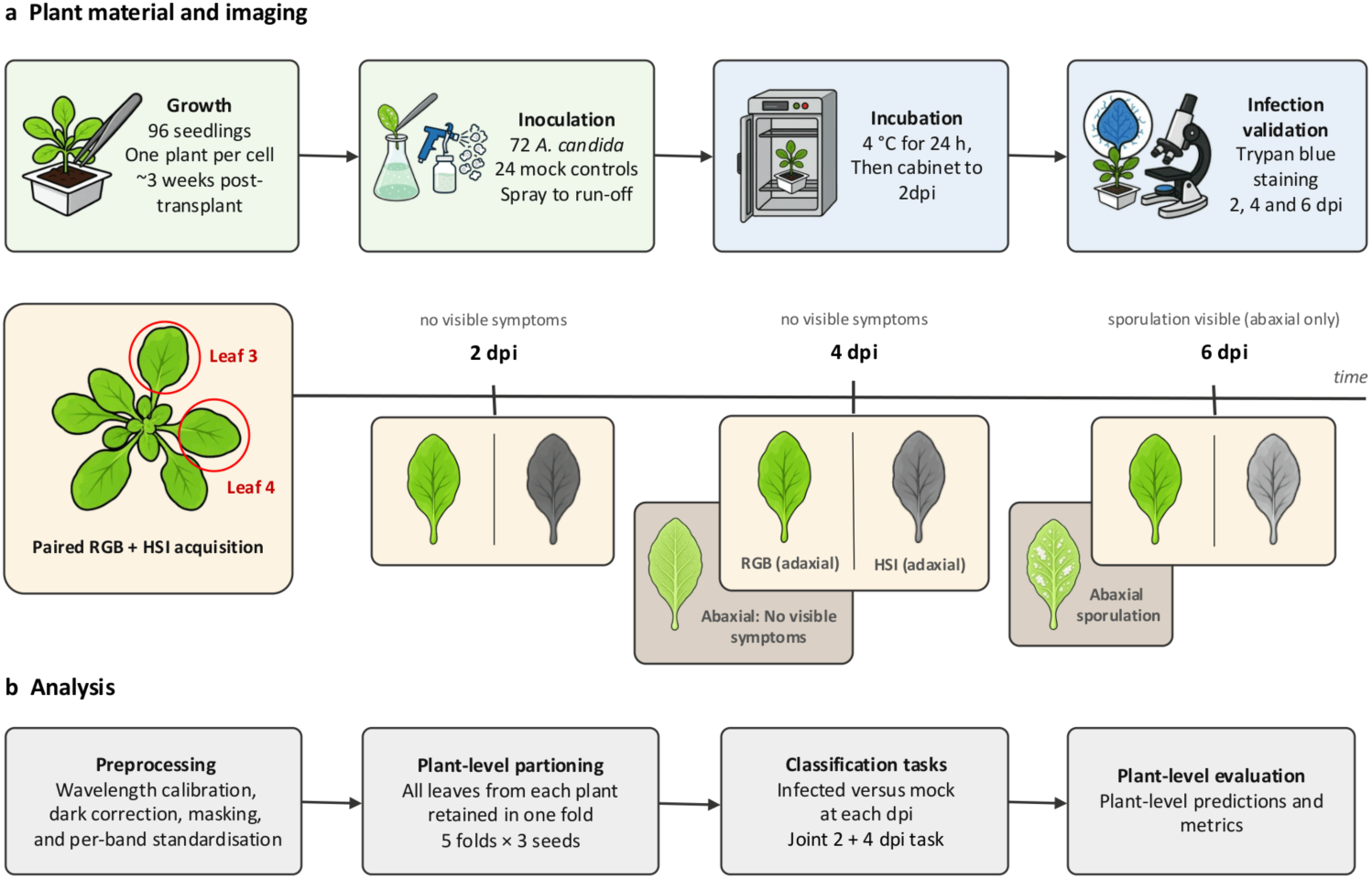
Sampling and plant-level evaluation workflow. The recovered inventory contains 96 plants and 576 paired observations. Archived multi-seed comparisons and recovered-cohort analyses are reported separately; treatment–tray and acquisition-order confounding remain.

**Figure 2.**
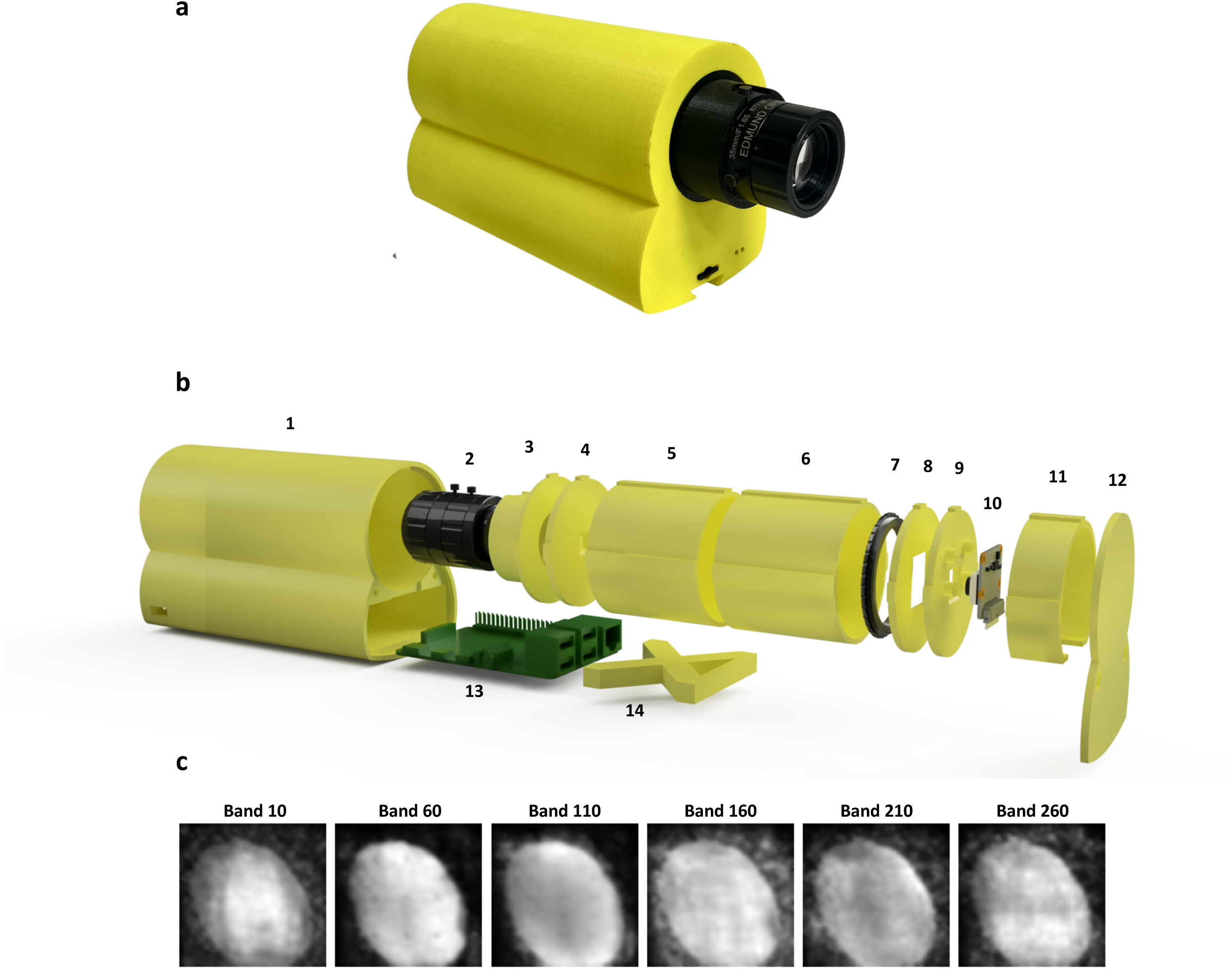
The low-cost hyperspectral imaging (HSI) unit. **(a)** Assembled unit (photograph). **(b)** Exploded view (CAD render) of the optical, structural and electronic components: 1 case; 2 35 mm Edmund Optics C Series VIS–NIR fixed-focal-length lens (67-716); 3 front lens; 4 square aperture; 5, 6 extension tubes; 7 +10 macro 52 mm lens; 8 diffraction grating; 9 camera base; 10 Raspberry Pi Camera Module 3 NoIR (12 MP); 11 seal extension; 12 lid; 13 Raspberry Pi 4 Model B; 14 X-axis extension. **(c)** Representative reconstructed hyperspectral images at bands 10, 60, 110, 160, 210 and 260.

The optical train uses a 35 mm Edmund Optics C Series VIS–NIR lens (part 67-716), the retained close-up lens and diffraction grating, and the existing enclosure (Fig. 2b). The original squareHSIBetaV1 software extracts the zero-order and first-order diffraction regions and assembles spectral cubes after wavelength calibration and dark correction [21]. Source images of 2592 × 1944 pixels were reconstructed into arrays of 313 × 132 × 135 spectral and spatial positions. These dimensions describe the reconstructed computational sampling. The acquisition size was retained for compatibility with the reconstruction software. Readout and exposure are controlled through libcamera on a Raspberry Pi 4B. Assembly and compatibility details are provided in Appendix S1.

The listed camera-unit components total approximately £489.20, including replacement prices for reused optical parts. The subtotal covers the camera unit and excludes printing, the laboratory illumination source, the copy stand and the reconstruction workstation. The cost breakdown is given in Appendix S1. Acquisition requirements are described in Section 2.3.

### 2.3. Acquisition set-up and protocol

All imaging was performed in a dark room to exclude ambient light. Both the HSI and RGB data were acquired top-down from the adaxial (upper) leaf surface only. Samples were illuminated with the Leica CLS 150 X source through dual fibre-optic arms placed symmetrically at approximately 45° to the sample plane. The camera was mounted on a copy stand about 10 cm above a matte-black background, and the source was warmed up for about 15 min at the start of each session and then left running continuously. Camera height and imaging geometry were kept consistent throughout a session and were not deliberately reset between trays. The external 35 mm lens was positioned for the required imaging geometry, and the Camera Module 3 autofocus was enabled in macro mode for fine focus, so focus was not fixed at infinity; live preview was used to position each sample. Images were captured with libcamera-still (JPEG quality 100, tungsten white-balance mode, cdn_hq denoising, 20 s preview, 2592 × 1944 pixels, macro autofocus range, RGB888 output). Exposure was fixed at 2 s (2,000,000 µs). Analogue gain was not set in the command and was under automatic control; it was not independently logged. Samples were processed sequentially and we tried to keep a similar acquisition order between sessions, but the order was not strictly fixed, formally randomised or logged. Inoculated and mock-inoculated plants were kept on separately marked trays and imaged with the same acquisition workflow. The mock tray was always imaged first and the inoculated tray second, at every session (2, 4 and 6 dpi), to reduce the risk of cross-contamination from infected material. Treatment was therefore consistently correlated with acquisition order, which is a potential acquisition-time confound.

Each plant was imaged at three time points. At each time point, leaves 3 and 4 of the rosette were imaged, with designated leaves followed across sessions and physically marked with a pipette marker. The design comprised 96 plants (72 inoculated and 24 mock-inoculated), two leaves and three time points, giving 576 paired HSI–RGB observations. The recovered inventory contains all 576 pairs, including 192 pairs at each time point. Plant identity was recorded using tray positions and leaf markers. The archived multi-seed classification analyses used the paired observations available when those runs were performed. Recovered-cohort results are explicitly identified below and do not retrospectively change the input population of the archived runs.

### 2.4. Spectral calibration and preprocessing

Spectral calibration followed Salazar-Vazquez and Mendez-Vazquez [21]. Fluorescent-lamp spectra, incandescent-lamp spectra and lens-covered dark frames were used to establish the wavelength mapping and apply dark correction in squareHSIBetaV1. The retained records do not establish whether dark frames were acquired separately for every tray. Each cube contains 313 computational spectral positions. Recovered MAT files and their matching hypercube headers contain identical nominal wavelength arrays spanning 400–1052.09 nm. The acquisition record identifies a narrower calibrated range of 400.000–968.487 nm covering 273 positions. The valid calibration range and correspondence with the historical training tensors have not been independently established for all stored positions; spectral tests are therefore reported by computational band index. No white reference panel was used, so these data are intensity measurements rather than absolute reflectance. Spectral resolution, calibration error, signal-to-noise ratio, sensor linearity and registration accuracy were not independently characterised (Table 1).

**Table 1.** Acquisition settings and instrument characteristics.

| Characteristic | Value | Status or note |
| --- | --- | --- |
| Illumination | Leica CLS 150 X, 150 W halogen, 3400 K; dual fibre-optic arms at $\sim 45^\circ$ | Excluded from the listed camera-component subtotal |
| Geometry and environment | Camera $\sim 10$ cm above sample on a copy stand; matte-black background; dark room; 15 min warm-up | Controlled conditions |
| Exposure | 2 s (2,000,000 μs), fixed | Fixed in the acquisition command |
| Analogue gain | Automatic (not set in the command) | Not independently logged; per-frame gain not verified |
| Acquisition command | libcamera-still: quality 100, tungsten white balance, cdn_hq denoising, 20 s preview, macro autofocus | Camera Module 3 autofocus enabled |
| Image and cube size | Source $2592 \times 1944$ px; cube $132 \times 135$ px | Below the native IMX708 resolution |
| Spectral positions | 313 computational positions; nominal wavelengths stored | 273 positions reported as calibrated; tensor correspondence unverified |
| Recorded wavelength ranges | Nominal 400–1052.09 nm; reported calibrated 400.000–968.487 nm | Nominal array is not evidence of calibration accuracy or full-range validity |
| Mean spacing of calibrated positions | $\approx 2.1$ nm | Sampling interval, not spectral resolution |

**Table 2.** Complete recovered inventory: 96 plants (72 inoculated and 24 mock-inoculated), with 192 paired observations at each dpi and 576 observations in total. These counts describe the recovered inventory. The four-class analysis is summarised separately in Table 5; supplementary archived evaluations retain their documented analysis populations.

| Task | Time point(s) (dpi) | Plants | Inoculated | Mock | Majority-class accuracy (%) | Includes 6 dpi (early symptomatic) |
| --- | --- | --- | --- | --- | --- | --- |
| 2 dpi | 2 | 96 | 72 | 24 | 75.00 | No |
| 4 dpi | 4 | 96 | 72 | 24 | 75.00 | No |
| 6 dpi | 6 | 96 | 72 | 24 | 75.00 | Yes |
| All dates | 2, 4, 6 | 96 | 72 | 24 | 75.00 | Yes |

The binary evaluation pipeline used HSI tensors with shape 313 × 132 × 135 and RGB images resized to 224 × 224. Band-wise HSI standardisation parameters were fitted only on the inner training plants of each outer fold and applied unchanged to validation and test plants. The available implementation does not verify the full upstream masking and reconstruction history of the archived tensors. For the recovered cohort, missing paired observations were appended while existing overlapping observations were retained without duplication. Processing equivalence between recovered files and archived tensors was not established; the combined-cohort results are interpreted with this limitation. The spectral analysis used the entire stored cube rather than a separately verified leaf mask.

### 2.5. Dataset and plant-level evaluation design

#### Tasks

The primary analysis uses the four-class labels: Healthy, 2 dpi, 4 dpi and 6 dpi. Healthy denotes the pooled mock-inoculated class, whereas the inoculated observations are labelled by imaging time point. Four-class accuracy is calculated from the diagonal of the aggregated confusion matrix. A derived binary result maps Healthy to the negative class and all three inoculated classes to the positive class in both the true and predicted labels. Supplementary binary analyses compare inoculated and mock-inoculated plants separately at 2, 4 and 6 dpi. We treat 2 and 4 dpi as presymptomatic because no sporulation was visible on either leaf surface at those times. At 6 dpi, sporulation was visible on the abaxial surface while the imaged adaxial surface remained symptom-free; this time point therefore provides an early symptomatic reference. The same plants contribute to the different time-point tasks.

#### Unit of analysis and splitting

For each supplementary binary model, task and training seed, predictions from the five outer test folds were pooled. The predicted inoculated-class probabilities of all available observations belonging to a plant were averaged; a mean probability ≥0.5 defined a positive prediction. AUROC was calculated from unthresholded plant-level probabilities. Outer folds were stratified by treatment at the plant level and reused across models and training seeds within each task; fold assignments were not identical across all tasks. For each outer training partition, 20% of its plants were reserved for inner validation, stratified by treatment with random state equal to the training seed plus the fold index. Checkpoints and early stopping were selected using inner-validation plant-level balanced accuracy, without using outer test outcomes.

#### Metrics

For the primary four-class analysis, accuracy is the sum of the diagonal counts divided by the total count in the confusion matrix. The derived binary accuracy is computed after collapsing both the true and predicted labels into healthy versus inoculated. These primary accuracies are reported separately from the supplementary plant-level binary metrics. For the supplementary binary analyses, in the complete design, 75% of plants were inoculated; predicting every plant as inoculated therefore gives 75% accuracy, 100% sensitivity, 0% specificity and 50% balanced accuracy. Balanced accuracy (BA, the mean of sensitivity and specificity) and AUROC are the primary metrics. Accuracy, F1, precision, sensitivity and specificity are also reported. Archived three-seed results are summarised by their arithmetic mean and sample SD over seeds 157, 257 and 357. This SD reflects training randomness, not uncertainty between experimental runs. Single-seed results have no between-seed SD. Table 8 retains the archived seed-157 plant-bootstrap intervals, computed from 5,000 treatment-stratified resamples of fixed out-of-fold predictions without retraining. For paired comparisons, we additionally resampled plants within treatment 5,000 times (bootstrap seed 20260929), using identical plant multiplicities for PSNet, its comparator and all three training seeds. We averaged the seed-specific differences in BA and AUROC within each replicate and used their 2.5th and 97.5th percentiles. These pointwise intervals are conditional on fitted models, are not adjusted for multiple comparisons, and do not constitute independent-run validation (Table S6).

#### Scope of the evaluation

Plant-disjoint folds drawn from one cultivation run share the growth cycle, inoculum preparation, calibration session, illumination set-up and operator. They guard against leakage between images of the same plant but are not independent of batch effects, and we do not present them as evidence of generalisation to other runs, sites or conditions.

### 2.6. Computational framework

#### 2.6.1 Overview

PSNet is a dual-stream network combining RGB and HSI features (Fig. 3). The RGB branch uses an ImageNet-pretrained ResNet34 [27, 28], global average pooling and a projection. The HSI branch uses the HSNet encoder (Fig. 4). The two embeddings are concatenated and passed to an MLP classifier. Figure 3 shows the four output categories of the primary task. The supplementary binary implementation uses 64-dimensional embeddings in each branch, a 128-dimensional concatenated vector and a two-output classifier. The following sections specify that binary implementation, its comparison models and training settings.

**Figure 3.**
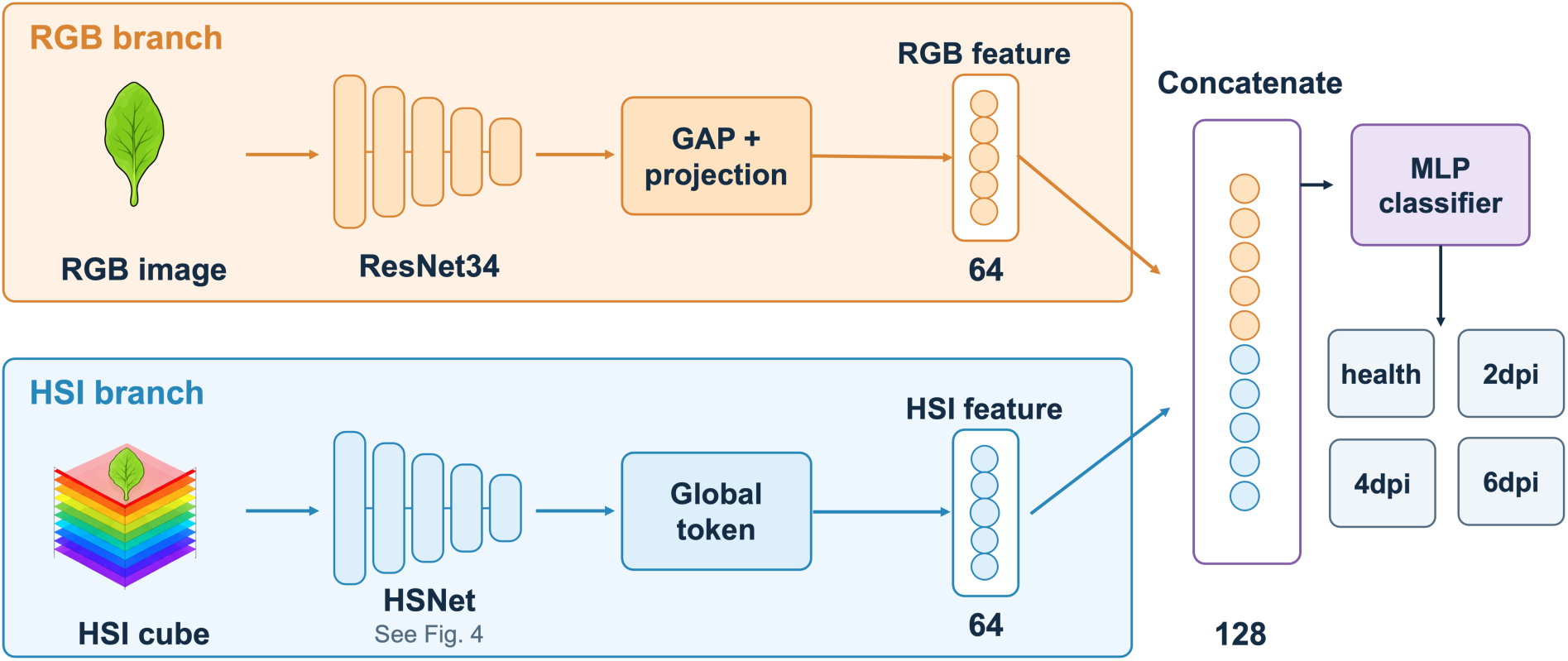
Overview of PSNet for the primary four-class task. RGB and hyperspectral inputs are encoded by the ResNet34 and HSNet branches, respectively. The resulting features are concatenated and passed to an MLP classifier with four output categories: Healthy, 2 dpi, 4 dpi and 6 dpi. For the supplementary date-specific binary evaluations, the classifier has two outputs representing mock-inoculated and inoculated plants.

**Figure 4.**
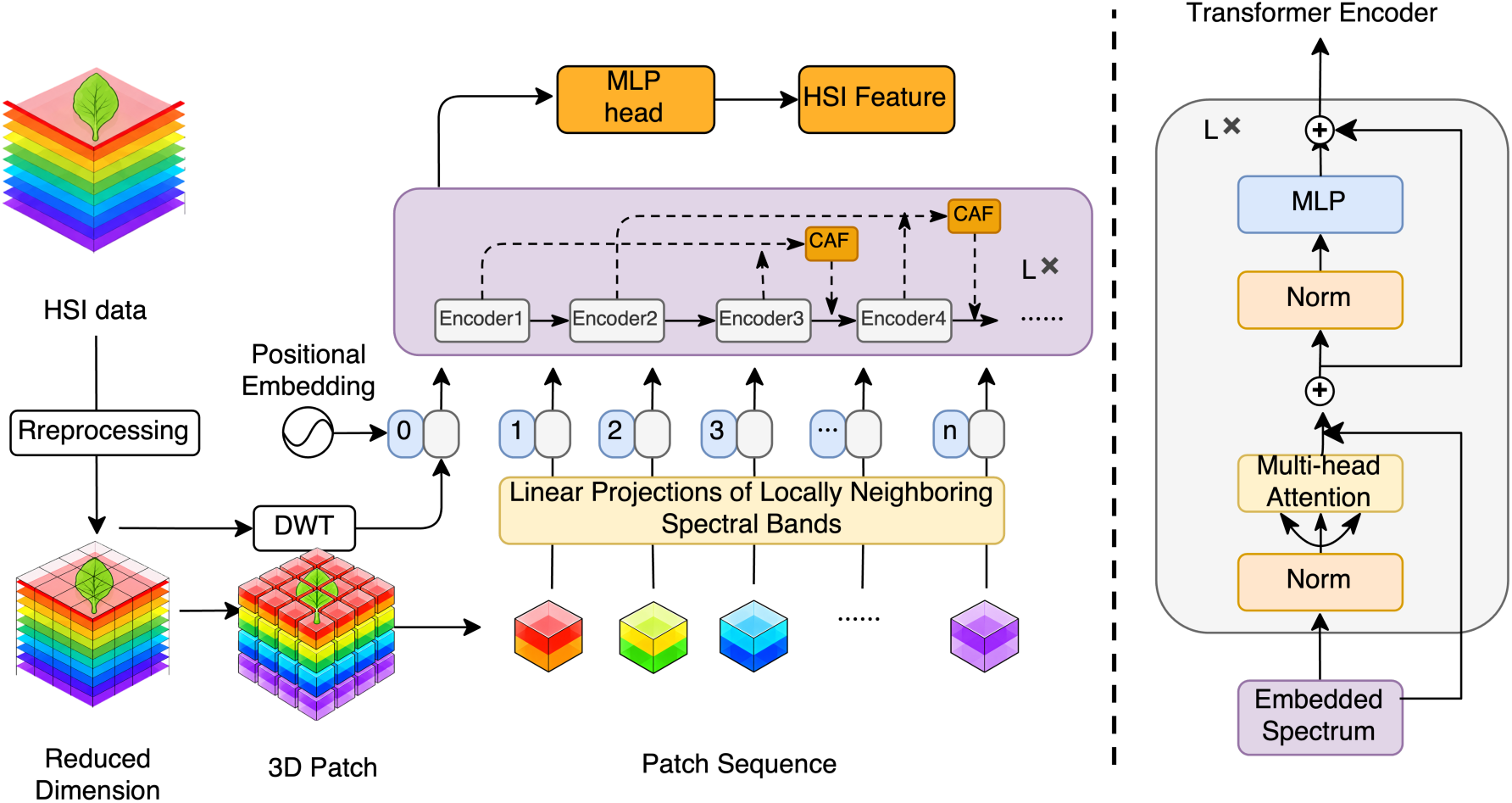
Architecture of the HSNet encoder. Left: a three-level Haar discrete wavelet transform (DWT) of the spatially averaged preprocessed input spectrum initialises the global token (token 0). In parallel, learned channel reduction and non-overlapping 3D patch embedding produce 512 patch tokens. Positional embeddings are added before the sequence is processed by five Transformer layers with cross-layer adaptive fusion (CAF). The final normalised global token provides the 64-dimensional HSI feature used by PSNet. Right: the structure of a Transformer layer, comprising layer normalisation, multi-head self-attention, an MLP and residual connections. L denotes the number of Transformer layers.

### 2.6.2 Spatial branch

The RGB branch is an ImageNet-pretrained ResNet34. The RGB image **X***_r_*_g*b*_ ∈ ℝ^224×224×3^ passes through the convolutional and residual stages, the final feature map is aggregated by global average pooling (GAP), and a projection head produces the RGB embedding **z***_r_*_g*b*_.

#### 2.6.3 Spectral branch

The spectral branch, HSNet, represents the hyperspectral cube as a sequence of tokens (Fig. 4). Let

**X**_ℎ*si*_ ∈ ℝ*^H^*^×W×*C*^ be the preprocessed cube with spatial size *H* × *W* and *C* spectral channels, and let

**x***_i_*_,j_ ∈ ℝ*^C^* be the spectrum at pixel (*i*, *j*). The input sequence consists of one global token and *N* patch tokens computed in parallel from the same cube.

Global token. The input cube is first averaged over its spatial dimensions to obtain a spectrum ̄x. A three-level Haar transform is then applied along its spectral axis. At each level, an odd-length input is extended by replicating its final value before pairwise sums and differences are computed:

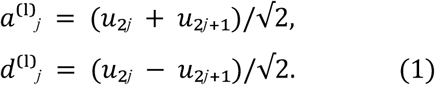

Here l = 1, 2, 3 indexes the decomposition levels; the first input is the mean spectrum ̄x;, and each subsequent input is the preceding approximation vector. The detail vectors are concatenated in level order, followed by the final approximation vector. The concatenation is truncated to the input band count C = 313:

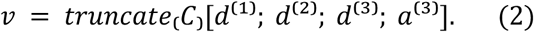

A learned linear projection with bias, followed by GELU, maps v into the 64-dimensional global token **x**_g_ . This token is sample-dependent, unlike a freely learned CLS vector. The operation is computed independently for each sample.

Patch tokens. A 3 × 3 two-dimensional convolution with padding 1 maps the 313 input bands to C′ = 64 learned feature channels, followed by batch normalisation and GELU. These are feature channels rather than retained physical wavelengths. Non-overlapping patches span 16 positions along each spatial axis and eight reduced channels. The number of patches along each axis is

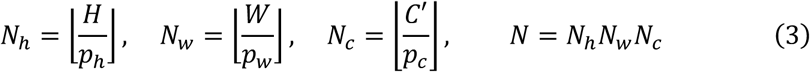

so that rows, columns or bands that do not fill a complete patch are discarded. The patch with index (*a*, *b*, *c*), where *a* ∈ {1, … , *N*_ℎ_}, *b* ∈ {1, … , *N*_w_} and *c* ∈ {1, … , *N_c_*}, is

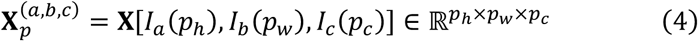

where *I_k_*(*p*) = {(*k* − 1)*p* + 1, … , *kp*} is the *k* th block of *p* consecutive indices. Each patch is embedded by a strided 3D convolution with kernel and stride (8, 16, 16) in channel-depth, height and width order. For an input of 64 × 132 × 135, the grid is 8 × 8 × 8, producing N = 512 patch tokens of dimension d = 64. Spatial remainders that do not fill a complete patch are discarded. The input sequence to the Transformer is

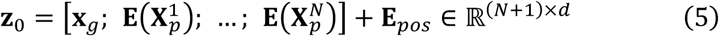

The first 513 positional embeddings are added to the global and patch tokens, followed by dropout. The registered position tensor contains 1,025 positions; the remaining 512 positions (32,768 elements) are unused for this input shape. Thus positional encoding participates in the forward pass, although not every registered element is used.

Encoder and cross-layer adaptive fusion (Fig. 4). Five Transformer layers use four attention heads each, with a feed-forward width of 256 and dropout 0.5. As shown on the right of Figure 4, each layer comprises pre-normalised multi-head self-attention and an MLP, both with residual connections. Cross-layer adaptive fusion (CAF), adapted from SpectralFormer [29], concatenates the current tokens with a cached earlier-layer input and applies a learned Linear(128, 64) projection. Writing hₗ for the tokens before optional fusion and uₗ for the actual input to Transformer layer l,

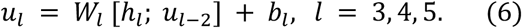

The first two layers use their incoming tokens without fusion. The cached inputs supplying layers 3, 4 and 5 are respectively those entering layers 1, 2 and 3. The projection is shared over token positions, and its output is passed to the corresponding Transformer layer. A final layer normalisation is applied, and the final global token is used as the HSI embedding.

#### 2.6.4 Fusion and classification

The RGB and HSI embeddings, each of dimension 64, are concatenated and classified by Linear(128, 128), GELU, dropout 0.5 and a final linear layer. The final layer has four outputs for the primary four-class task and two outputs for the supplementary binary implementation. Training minimises the unweighted cross-entropy loss:

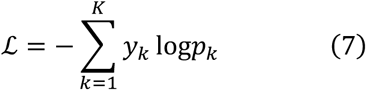

Here K is the number of output classes: K = 4 for the primary four-class task and K = 2 for the supplementary binary implementation; *y_k_* is the one-hot label and *p_k_* is the predicted probability. The supplementary implementation uses ordinary shuffled training batches without class-weighted loss or class-balanced resampling.

#### 2.6.5 Baselines and ablation variants

All supplementary binary model comparisons within a task use the same outer plant folds and training seeds. RGB-only baselines are ImageNet-pretrained ResNet18 and ResNet34 with 64-dimensional projections and two-class linear heads. The HSI-only Transformer uses the same HSI encoder as PSNet followed by a linear classifier. The spectral 1D CNN takes the spatially averaged 313-position spectrum and applies three 1D convolutional blocks and global pooling. The compact 3D CNN takes the full cube through three strided 3D convolutional blocks and global pooling. The simple multimodal model combines a ResNet18 embedding with the spectral 1D CNN embedding using an MLP; the plain multimodal model uses a learned CLS token with CAF disabled. Registered trainable parameter counts are 11,209,474 for RGB ResNet18, 21,317,634 for RGB ResNet34, 672,258 for the HSI Transformer, 23,234 for the spectral 1D CNN, 81,474 for the compact 3D CNN and 11,240,834 for the simple multimodal model. PSNet variants are defined in Table 3.

**Table 3.** PSNet variants and registered trainable parameter counts. Counts include 24,768 bypassed fusion parameters when CAF is disabled and 32,768 unused positional-embedding elements in the full 3D path. Equal registered counts do not establish equal active capacity. Baseline definitions and parameter counts are given in Section 2.6.5.

| Configuration | Global token | Patch tokens | Registered parameters | Change relative to full PSNet |
| --- | --- | --- | --- | --- |
| Full PSNet | Wavelet-initialised | 512 | 22,006,402 | — |
| CAF disabled | Wavelet-initialised | 512 | 22,006,402 | Cross-layer adaptive fusion bypassed |
| Mean patch token | Mean of patch tokens | 512 | 21,986,306 | Wavelet-initialised token replaced by mean of patch tokens |
| Alternative 2D tokenisation | Wavelet-initialised | 64 | 26,823,042 | 2D tokenisation (changes reduction stage, token number and capacity) |
| Learnable CLS | Learnable parameter | 512 | 21,986,370 | Wavelet-initialised token replaced by learnable CLS token |
| Plain multimodal | Learnable parameter | 512 | 21,986,370 | Learnable CLS token and CAF disabled |

#### 2.6.6 Training and reproducibility

Models were trained end-to-end with AdamW [30] and unweighted cross-entropy. RGB and HSI inputs received synchronised horizontal and vertical flips, each independently with probability 0.5. The archived configurations record embedding dimension 64, five Transformer layers, four attention heads, dropout 0.5 and pretrained RGB backbones. Inner-validation plant-level BA selected checkpoints and governed early stopping. The retained implementation contains no label smoothing, stochastic depth, learning-rate warm-up or ReduceLROnPlateau schedule. The documented run protocol specifies learning rate 3 × 10⁻⁴, weight decay 10⁻⁴, batch size 8, at most 100 epochs and early-stopping patience 12. These are protocol settings; the older run manifests do not independently record every optimisation setting. No auditable model-specific hyperparameter-search history was available, so equivalent tuning effort is not claimed. Archived runs used an NVIDIA A100-SXM4-80GB GPU. Code is identified under Data availability.

### 2.7 Spectral statistics

For each stored HSI cube, intensities were averaged across all spatial pixels, and available leaf spectra were then averaged within each plant and time point. The recovered inventory provided 72 inoculated and 24 mock plants at every time point. We compared the two treatments separately at each of the 313 computational spectral positions using two-sided Welch tests, with Hedges’ g defined for inoculated minus mock plants. Benjamini–Hochberg correction [31] was applied separately within each dpi across its 313 tests; q < 0.05 defined the primary significant-band count. A secondary correction across all 939 tests is reported separately. The minimum q in Table 9 refers to the within-dpi correction. This analysis uses whole-cube intensity averages, not separately masked leaf tissue or calibrated reflectance. Bootstrap intervals for spectral effect sizes, physical red-edge metrics and models adjusting for exposure, gain or background were not estimated in this analysis; the required calibration and acquisition covariates were not verified.

## 3 Results

### 3.1 Dataset, symptom status and infection validation

No visible symptoms were present at 2 or 4 dpi. Images were acquired from the adaxial surface. At 6 dpi, white-blister sporulation was visible on the abaxial surface while the imaged adaxial surface remained visually symptom-free (Fig. 5). Thus, 6 dpi represents an early symptomatic reference rather than evidence of whole-plant presymptomatic detection. The recovered dataset contains 96 plants and 192 paired observations at every time point, with 72 inoculated and 24 mock plants (Table 2). The primary four-class results are reported from the confusion matrix. Supplementary multi-seed binary estimates retain their documented evaluation scope, while the spectral analyses use the recovered inventory. Qualitative staining does not establish an infection rate for every imaged plant.

**Figure 5.**
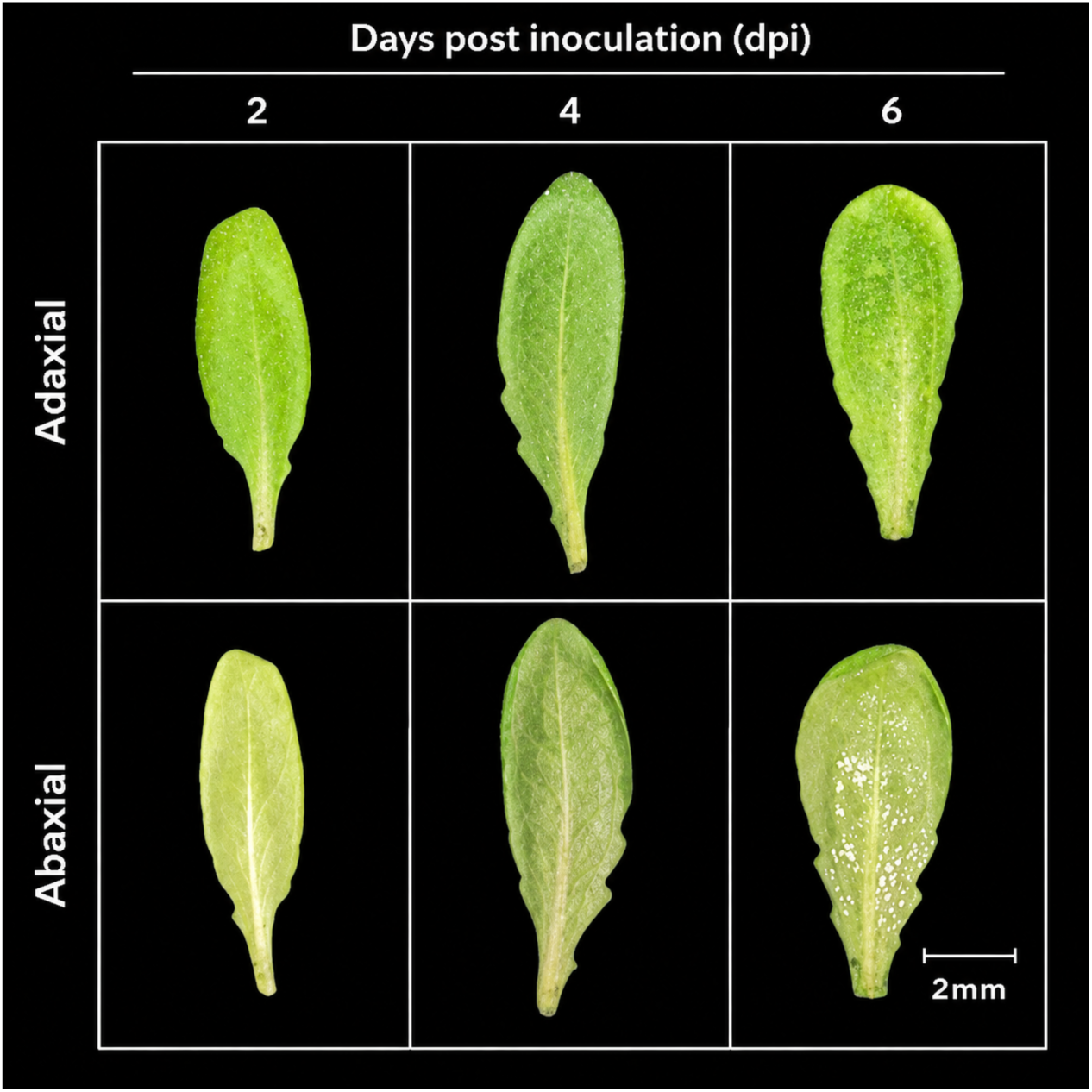
Representative adaxial and abaxial surfaces of leaves at 2, 4 and 6 days post inoculation (dpi) with *Albugo candida*. No symptoms are visible at 2 or 4 dpi; white sporulation is visible on the abaxial surface at 6 dpi, whereas the adaxial surface remains symptom-free. Scale bar, 2 mm. Images illustrate symptom status; they do not provide a quantified treatment comparison.

### 3.2 Instrument characterisation

Table 1 lists the instrument characteristics that were available at the time of writing. Spectral resolution, wavelength calibration error, signal-to-noise ratio at the final exposure, spatial resolution, repeatability, sensor linearity and RGB–HSI registration accuracy have not yet been measured or verified, and we therefore make no claims about them. Preliminary exposure tests at 10–100 ms did not use the final 2 s setting and are not sufficient to establish signal-to-noise ratio or the absence of saturation.

### 3.3 Classification performance

On the four-class task comprising Healthy, 2 dpi, 4 dpi and 6 dpi, PSNet achieved an overall accuracy of 90.00 ± 2.34%, precision of 90.35 ± 2.58%, an F1-score of 0.8996 ± 0.0239 and an averaged AUROC of 0.9878 ± 0.0025 (Table 5). In the aggregated confusion matrix, 288 of 320 predictions were correct. Class-wise recall was 95.00% for Healthy, 83.75% for 2 dpi, 87.50% for 4 dpi and 93.75% for 6 dpi, with the largest off-diagonal counts occurring between the 2 and 4 dpi categories.

**Table 4.** Tensor shapes and documented training protocol. Architecture settings are recorded in the archived configurations. Optimisation settings describe the retained run protocol; the older manifests do not independently capture every runtime setting.

| Item | Setting |
| --- | --- |
| RGB input | $224 \times 224 \times 3$ |
| HSI input after preprocessing | $132 \times 135$ spatial × 313 input bands; 64 learned channels in the patch branch |
| 3D patch size ($p_c \times p_h \times p_w$) | 8 channel-depth × 16 height × 16 width |
| Tokens | 1 global token + 512 patch tokens |
| Embedding dimension $d$ | 64 |
| Transformer layers; attention heads | 5 layers; 4 attention heads |
| Classification head | $128 \rightarrow 128 \rightarrow 4$ (primary); $128 \rightarrow 128 \rightarrow 2$ (supplementary binary); GELU and dropout 0.5 |
| Optimiser | AdamW; learning rate $3 \times 10^{-4}$; weight decay $10^{-4}$ (documented protocol) |
| Batch size; epochs | 8; maximum 100 epochs; patience 12 (documented protocol) |
| Learning-rate schedule | No warm-up or scheduler in the retained implementation |
| Loss | Unweighted cross-entropy; no class-balanced resampling |
| Random seeds | 157, 257, 357 for archived comparisons; 157 for the recovered pooled run |
| Augmentation and regularisation | Synchronous horizontal/vertical flips ($p = 0.5$ each); dropout 0.5; inner-validation early stopping |
| Training hardware | NVIDIA A100-SXM4-80GB for archived runs |

**Table 5.** Archived PSNet classification performance. Mean ± SD across seeds 157, 257 and 357, each evaluated with five plant-disjoint folds and available observations aggregated within plants. These estimates, including the standalone 2 dpi values, are retained from the archived analysis and are not recalculated recovered-cohort estimates. BA is balanced accuracy.

| Task | Accuracy (%) | BA (%) | AUROC (%) | F1 (%) | Precision (%) | Sensitivity (%) | Specificity (%) |
| --- | --- | --- | --- | --- | --- | --- | --- |
| Four-class | $90.00 \pm 2.34$ | - | $98.78 \pm 0.25$ | $89.96 \pm 2.39$ | $90.35 \pm 2.58$ | - | - |
| 2 dpi | $69.79 \pm 3.61$ | $55.79 \pm 4.01$ | $60.46 \pm 4.90$ | $80.55 \pm 3.80$ | $77.12 \pm 2.03$ | $84.54 \pm 7.98$ | $27.78 \pm 10.49$ |
| 4 dpi | $88.54 \pm 2.08$ | $83.56 \pm 4.24$ | $93.88 \pm 2.60$ | $92.45 \pm 1.30$ | $91.46 \pm 2.61$ | $93.52 \pm 0.80$ | $73.61 \pm 8.67$ |
| 6 dpi | $95.14 \pm 1.20$ | $95.37 \pm 0.80$ | $98.99 \pm 0.60$ | $96.68 \pm 0.89$ | $98.60 \pm 1.39$ | $94.91 \pm 2.89$ | $95.83 \pm 4.17$ |

Collapsing the same true and predicted labels into healthy versus inoculated yielded 97.50% binary accuracy, corresponding to 312 correct predictions out of 320. This derived result summarises the four-class predictions across all dates, with Healthy as the negative class and the three inoculated time-point classes combined as the positive class.

Supplementary binary analyses compared inoculated and mock-inoculated plants separately at each imaging time point using plant-level probability aggregation. Balanced accuracy was 55.79 ± 4.01% at 2 dpi, 83.56 ± 4.24% at 4 dpi and 95.37 ± 0.80% at 6 dpi. The 2 dpi balanced accuracy was calculated from the three seed-specific plant-level confusion matrices in Table S5. At 4 dpi, before visible symptoms, PSNet achieved an AUROC of 0.939 ± 0.026, with sensitivity of 93.52 ± 0.80% and specificity of 73.61 ± 8.67%. At 6 dpi, AUROC was 0.990 ± 0.006; abaxial sporulation was present even though the imaged adaxial surface remained symptom-free. The four-class and derived binary accuracies provide the primary overall classification results, while the supplementary analyses report discrimination within individual imaging dates.

### 3.4 Comparison with RGB-only, HSI-only and multimodal baselines

Model rankings changed markedly with time after inoculation (Table 6, Fig. 6; full metrics in Tables S1–S3). Four observations stand out. First, no model that received RGB input exceeded a balanced accuracy of 63.4% at 2 dpi (range 50.7–63.4%; AUROC 0.509–0.711). Second, at 4 dpi every model with an RGB stream, including RGB-only ResNets, reached balanced accuracies of 83.6–92.4% (AUROC 0.930–0.974); PSNet had the lowest balanced accuracy in this group (83.56%), and the plain multimodal model the highest (92.36%), with an RGB-only ResNet18 close behind (90.28%). Third, at 6 dpi the same models reached 91.2–97.5% balanced accuracy; RGB ResNet18 (95.14%) fell within the range of the multimodal models (93.29–97.45%), whereas RGB ResNet34 (91.20%) was below it. Fourth, the HSI-only models behaved differently from all others. The HSI Transformer and the compact 3D CNN were the only models to discriminate well at 2 dpi (balanced accuracy 76.15% and 86.05%; AUROC 0.850 and 0.916), yet at 4 dpi their balanced accuracy fell to 58.80% and 58.56%, and at 6 dpi to 54.17% and 50.00%. At 6 dpi the compact 3D CNN labelled every plant as infected in all three seeds. The one-dimensional spectral CNN was at or near chance at every time point (balanced accuracy 54.80%, 50.00% and 50.23% at 2, 4 and 6 dpi).

**Figure 6.**
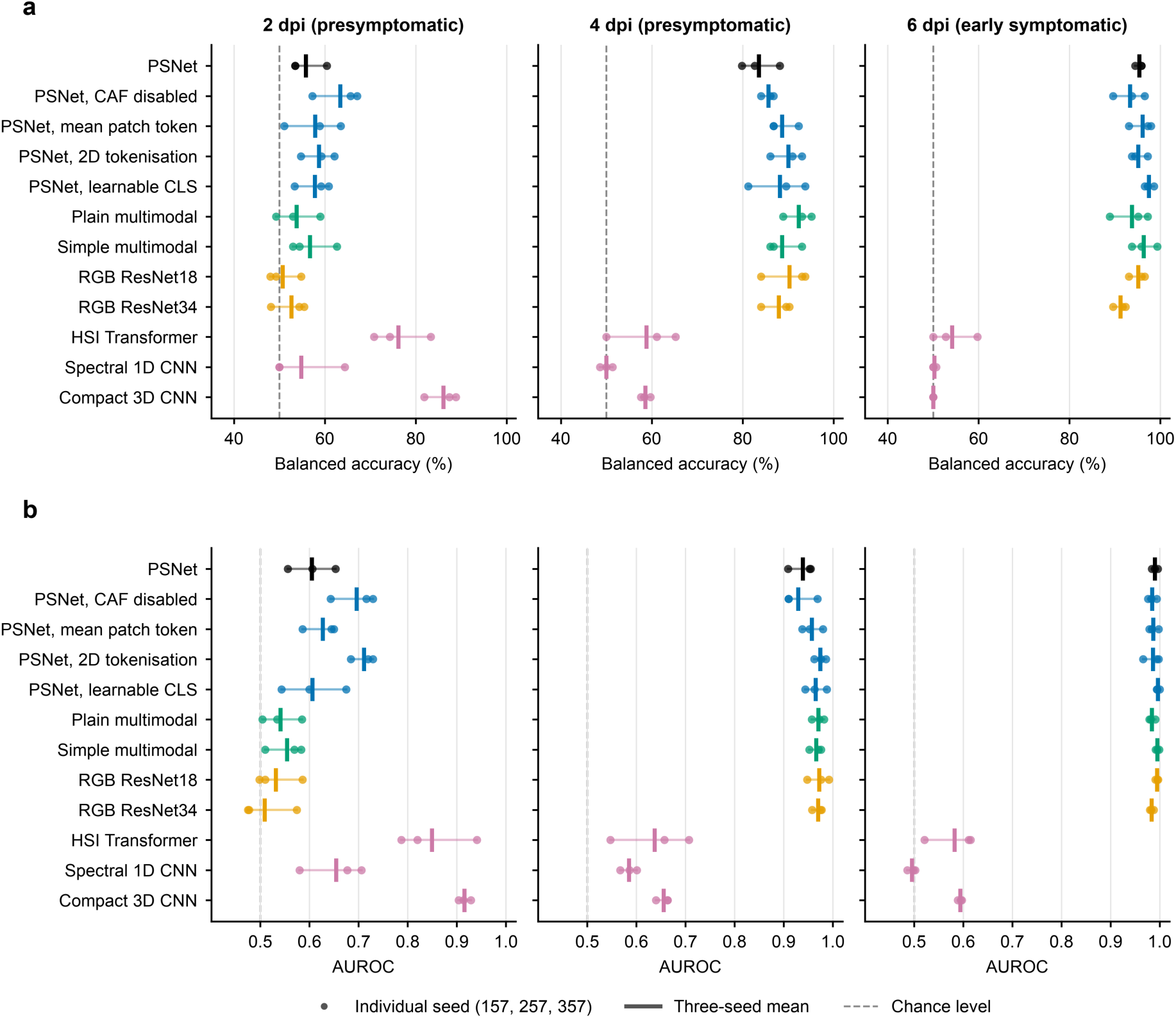
Supplementary date-specific classification performance. (a) Balanced accuracy and (b) AUROC for inoculated versus mock-inoculated classification at 2, 4 and 6 days post inoculation. Dots show individual seeds (157, 257 and 357), vertical marks show their arithmetic mean, and horizontal lines connect the seed values. The dashed line indicates chance level. The PSNet 2 dpi BA values are calculated from the counts in Table S5; AUROC and the other plotted estimates retain the archived prediction-based evaluation scope.

**Table 6.** Balanced accuracy (BA) and AUROC of all models at 2, 4 and 6 dpi. Mean ± SD across three seeds, each with five plant-disjoint folds. These are archived available-observation estimates. Paired pointwise bootstrap intervals are in Table S6. All metrics are in Tables S1–S3.

| Model | 2 dpi BA (%) | 2 dpi AUROC | 4 dpi BA (%) | 4 dpi AUROC | 6 dpi BA (%) | 6 dpi AUROC |
| --- | --- | --- | --- | --- | --- | --- |
| PSNet | 55.79 ± 4.01 | 0.605 ± 0.049 | 83.56 ± 4.24 | 0.939 ± 0.026 | 95.37 ± 0.80 | 0.990 ± 0.006 |
| CAF disabled | 63.35 ± 5.33 | 0.696 ± 0.046 | 85.65 ± 1.45 | 0.930 ± 0.034 | 93.29 ± 3.50 | 0.985 ± 0.009 |
| Mean patch token (replaces wavelet token) | 57.82 ± 6.27 | 0.627 ± 0.035 | 88.66 ± 3.21 | 0.957 ± 0.021 | 96.06 ± 2.63 | 0.987 ± 0.010 |
| Alternative 2D tokenisation | 58.70 ± 3.74 | 0.711 ± 0.024 | 90.05 ± 3.56 | 0.974 ± 0.012 | 95.14 ± 1.84 | 0.986 ± 0.017 |
| Learnable CLS token | 57.79 ± 3.94 | 0.606 ± 0.066 | 88.19 ± 6.36 | 0.965 ± 0.022 | 97.45 ± 1.06 | 0.996 ± 0.003 |
| Plain multimodal | 53.74 ± 4.89 | 0.541 ± 0.041 | 92.36 ± 3.18 | 0.971 ± 0.013 | 93.75 ± 4.34 | 0.984 ± 0.007 |
| Simple multimodal | 56.70 ± 5.23 | 0.554 ± 0.038 | 88.66 ± 3.82 | 0.966 ± 0.013 | 96.30 ± 2.81 | 0.996 ± 0.004 |
| RGB ResNet18 | 50.69 ± 3.61 | 0.531 ± 0.047 | 90.28 ± 5.42 | 0.972 ± 0.022 | 95.14 ± 1.84 | 0.994 ± 0.003 |
| RGB ResNet34 | 52.63 ± 3.96 | 0.509 ± 0.057 | 87.96 ± 3.43 | 0.970 ± 0.011 | 91.20 ± 1.45 | 0.984 ± 0.004 |
| HSI Transformer | 76.15 ± 6.46 | 0.850 ± 0.081 | 58.80 ± 7.90 | 0.637 ± 0.081 | 54.17 ± 5.01 | 0.582 ± 0.053 |
| Spectral 1D CNN | 54.80 ± 8.32 | 0.654 ± 0.066 | 50.00 ± 1.39 | 0.585 ± 0.017 | 50.23 ± 0.40 | 0.496 ± 0.009 |
| Compact 3D CNN | 86.05 ± 3.68 | 0.916 ± 0.012 | 58.56 ± 1.06 | 0.656 ± 0.014 | 50.00 ± 0.00 | 0.594 ± 0.004 |

Paired plant-bootstrap comparisons of the archived three-seed estimates are reported in Table S6. At 4 dpi, PSNet minus RGB ResNet18 was −6.71 percentage points in BA (pointwise 95% interval −12.96 to −0.69) and −0.0334 in AUROC (−0.0667 to −0.0060). Relative to the plain multimodal model, the corresponding differences were −8.80 percentage points (−15.05 to −3.24) and −0.0318 (−0.0631 to−0.0066). The comparisons use matched plant-level out-of-fold predictions and the conditional bootstrap procedure described in Section 2.5.

### 3.5 Contribution of individual PSNet components

Table 7 compares PSNet with four variants for which complete three-seed results are available. Variant rankings differed across the three imaging dates. At 6 dpi, balanced accuracy was 95.37% for PSNet, 93.29% with CAF disabled, 95.14% with alternative 2D tokenisation, 96.06% with a mean patch token and 97.45% with a learnable CLS token. The table reports the mean and between-seed SD for each configuration. Parameter counts (Table 3) were similar for all variants except alternative 2D tokenisation (26.8 million versus 22.0 million).

**Table 7.**
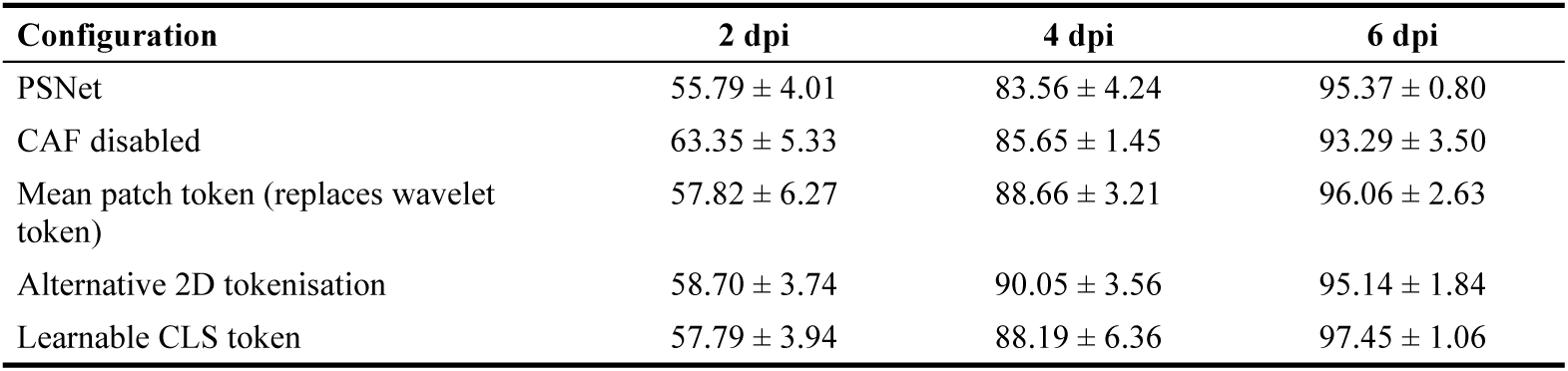
Balanced accuracy (%) of PSNet and variants with three complete seeds. Mean ± SD.

**Table 8.** Archived PSNet seed-157 plant-bootstrap intervals from 5,000 treatment-stratified resamples of fixed out-of-fold predictions without retraining. These intervals describe the archived single-date available-observation runs.

| Task | Accuracy (%) | BA (%) | AUROC | F1 |
| --- | --- | --- | --- | --- |
| 2 dpi | 55.91–73.12 | 42.57–63.50 | 0.423–0.687 | 0.692–0.823 |
| 4 dpi | 82.29–93.75 | 72.92–91.67 | 0.913–0.986 | 0.884–0.959 |
| 6 dpi | 91.67–98.96 | 90.28–99.31 | 0.973–1.000 | 0.942–0.993 |

**Table 9.** Recovered-cohort plant-level whole-cube spectral statistics. Each dpi includes 72 inoculated and 24 mock plants, with 313 valid Welch tests. Significance and minimum q use BH correction separately within each dpi. Hedges’ g is computed for inoculated minus mock; the final column summarises its absolute magnitude across spectral positions, without a bootstrap interval.

| Time point | Plants<br>(inoculated / mock) | Bands tested | Significant bands<br>(within-dpi BH) | Minimum q | Median \|Hedges’ g\| |
| --- | --- | --- | --- | --- | --- |
| 2 dpi | 72 / 24 | 313 | 269 | 0.001523 | 0.661280 |
| 4 dpi | 72 / 24 | 313 | 0 | 0.089157 | 0.444105 |
| 6 dpi | 72 / 24 | 313 | 0 | 0.170346 | 0.158316 |

Four further controlled substitutions (a raw-spectrum global token, current-layer input to CAF, their combination and fixed depth averaging) were specified but have not been run on the real data, so no conclusion is drawn about them.

### 3.6 Uncertainty of the estimates

Plant-bootstrap intervals for PSNet (seed 157) are reported in Table 8. At 2 dpi, the intervals were 42.6–63.5% for balanced accuracy and 0.423–0.687 for AUROC. At 4 dpi, they were 72.9–91.7% for balanced accuracy and 0.913–0.986 for AUROC. At 6 dpi, the lower bound of balanced accuracy was 90.3%. These intervals describe sampling variability of plants within this run, conditional on the fitted models. Raw plant-level confusion counts for each seed are given in Table S5.

### 3.7 Within-time-point spectral differences

In the recovered inventory, all 313 spectral positions were tested at each dpi using plant-level whole-cube mean spectra (Table 9). With BH correction applied separately within each dpi, 269 positions were significant at 2 dpi, whereas none met q < 0.05 at 4 or 6 dpi. Minimum within-dpi q values were 0.001523, 0.089157 and 0.170346, respectively. Median absolute Hedges’ g was 0.661280 at 2 dpi, 0.444105 at 4 dpi and 0.158316 at 6 dpi. A separate BH correction over all 939 tests yielded 213, 12 and 23 significant positions, respectively; these counts use a different testing family and are not substituted for the within-dpi results. Non-significance does not establish absence of spectral differences. The broad 2 dpi separation could reflect biological or acquisition-related structure, and whole-cube averages, non-reflectance intensities and processing differences limit physiological interpretation.

Table 9. Recovered-cohort plant-level whole-cube spectral statistics. Each dpi includes 72 inoculated and 24 mock plants, with 313 valid Welch tests. Significance and minimum q use BH correction separately within each dpi. Hedges’ g is computed for inoculated minus mock; the final column summarises its absolute magnitude across spectral positions, without a bootstrap interval.

### 3.8 Scope of model attribution

Archived band-group occlusion records showed unchanged BA and AUROC, but the corresponding masked probabilities were not retained. Whole-HSI and whole-RGB interventions were not validated. These records therefore cannot establish whether PSNet used a particular modality or spectral position, and no mechanistic attribution is claimed. The within-dpi spectral statistics describe treatment-group differences rather than learned model reliance.

### 3.9 Computational cost

Full PSNet has 22,006,402 registered trainable parameters (Table 3). Model-only inference on an NVIDIA A100-SXM4-80GB GPU, with batch size 1 and synthetic inputs, took 4.70 ms with peak GPU memory of 142.8 MiB. This excludes acquisition, reconstruction and preprocessing. Comparable latency measurements for all baselines, validated FLOP counts and end-to-end timings on the target hardware were not available; the GPU benchmark does not establish Raspberry Pi or edge-device performance.

## 4 Discussion

### 4.1 Principal findings

The primary finding is that PSNet classified Healthy, 2 dpi, 4 dpi and 6 dpi observations with 90.00% overall accuracy in the four-class evaluation. Collapsing these predictions into healthy versus inoculated labels yielded 97.50% accuracy across all dates. These results document the classification performance of the combined RGB–HSI framework under controlled experimental conditions. Supplementary date-specific binary analyses assessed discrimination using controls from the same imaging time point. At 4 dpi, before visible symptoms, PSNet achieved 83.56 ± 4.24% balanced accuracy and an AUROC of 0.939 ± 0.026. The primary analysis describes classification across the four categories, while the supplementary analysis supplies the time-specific evidence for presymptomatic classification.

### 4.2 Imaging time and modality performance

The supplementary modality comparisons showed different temporal patterns. At 2 dpi, the HSI Transformer and compact 3D CNN achieved AUROCs of 0.850 and 0.916, respectively, while PSNet had a balanced accuracy of 55.79 ± 4.01%. The recovered-cohort whole-cube spectral analysis also showed its broadest within-date group separation at 2 dpi. At 4 and 6 dpi, models receiving RGB input performed more strongly. At 4 dpi, RGB ResNet18 and the plain multimodal model achieved balanced accuracies of 90.28% and 92.36%, respectively, compared with 83.56% for PSNet. These patterns make imaging time an important consideration when selecting modalities for this task.

RGB classification at 4 and 6 dpi may draw on subtle differences in leaf colour, texture or posture, because the imaged surface showed no visible sporulation. The different temporal pattern of HSI may reflect changes in the information captured by spectral imaging. These are possible explanations of the observed model behaviour and remain hypotheses. The comparisons provide a practical basis for evaluating RGB-only and multimodal configurations at each imaging time point.

### 4.3 Architectural configuration

The ablation study examined how PSNet performance changed with the fusion mechanism, global token and tokenisation scheme. Variant rankings differed across imaging dates. At 6 dpi, PSNet achieved 95.37% balanced accuracy, compared with 93.29% with CAF disabled, 95.14% with alternative 2D tokenisation, 96.06% with a mean patch token and 97.45% with a learnable CLS token. These results describe the sensitivity of the classifier to its architectural configuration and show that component selection should be considered together with the imaging time point. PSNet combines the evaluated design elements in a single multimodal framework, while the variants provide reference configurations for this dataset.

### 4.4 Applications and future work

The compact, repairable imaging unit, USB-C power supply and open design files make the platform accessible for controlled-environment plant phenotyping. Pairing the unit with RGB imaging and plant-level model evaluation provides a workflow for comparing sensing modalities and tracking classification performance across infection stages. The hardware design and acquisition details are documented in the supplementary material, allowing the main text to focus on the biological task and model evaluation.

Future work will focus on expanding the dataset through additional experimental runs and larger plant cohorts, and on evaluating a broader range of pathogens and plant–pathogen systems. These extensions would assess the reproducibility of the observed temporal patterns and the transferability of the imaging and classification workflow.

## 5 Conclusions

This study presented PSNet, a dual-stream RGB–HSI classification framework, together with a low-cost hyperspectral imaging unit for the *Arabidopsis thaliana*–*Albugo candida* system. The primary four-class evaluation achieved 90.00% overall accuracy, and collapsing the same predictions into healthy versus inoculated labels yielded 97.50% accuracy across all dates. Supplementary date-specific binary evaluation achieved 83.56 ± 4.24% balanced accuracy and an AUROC of 0.939 ± 0.026 at 4 dpi, before visible symptoms. Additional modality and architectural comparisons characterised performance across imaging dates. Together, the primary classification results and supplementary analyses provide a basis for evaluating low-cost multimodal plant imaging under controlled conditions.

## Authorship contribution statement

**Gabriel U. Crabb**: Methodology (imaging system and experimental workflow), Investigation, Data Curation, Formal Analysis (exploratory modelling), Writing - Review & Editing. **Yiding Zhao**: Software, Methodology (PSNet architecture and training and validation framework), Writing - Review & Editing. **Xi Chen**: Supervision, Methodology (machine learning and computational framework), Writing - Review & Editing. **Nicholas K. Priest**: Supervision, Writing - Review & Editing. **Volkan Cevik**: Conceptualization, Supervision, Writing - Review & Editing.

## Acknowledgements

The authors thank the CFFF platform of Fudan University for providing computational resources and the University of Bath for supporting this research.

## References

1. Savary, S., et al., The global burden of pathogens and pests on major food crops. Nature Ecology & Evolution, 2019. 3(3): p. 430–439.

2. FAO, The impact of disasters and crises on agriculture and food security: 2021. 2021: Rome.

3. Hossain, M.M., et al., Plant disease dynamics in a changing climate: impacts, molecular mechanisms, and climate-informed strategies for sustainable management. Discover Agriculture, 2024. 2(1): p. 132.

4. Ristaino, J.B., et al., The persistent threat of emerging plant disease pandemics to global food security. Proceedings of the National Academy of Sciences, 2021. 118(23): p. e2022239118.

5. Fisher, M.C., et al., Worldwide emergence of resistance to antifungal drugs challenges human health and food security. Science, 2018. 360(6390): p. 739–742.

6. Lucas, J.A., N.J. Hawkins, and B.A. Fraaije, The evolution of fungicide resistance. Advances in Applied Microbiology , 2015. 90: p. 29–92.

7. Mahlein, A.K., Plant Disease Detection by Imaging Sensors - Parallels and Specific Demands for Precision Agriculture and Plant Phenotyping. Plant Dis, 2016. 100(2): p. 241–251.

8. Lau, H.Y. and J.R. Botella, Advanced DNA-Based Point-of-Care Diagnostic Methods for Plant Diseases Detection. Frontiers in Plant Science, 2017. Volume 8 - 2017.

9. Notomi, T., et al., Loop-mediated isothermal amplification (LAMP): principle, features, and future prospects. J Microbiol, 2015. 53(1): p. 1–5.

10. Hak, H., et al., Rapid on-site detection of crop RNA viruses using CRISPR/Cas13a. Journal of Experimental Botany, 2024.

11. Lowe, A., N. Harrison, and A.P. French, Hyperspectral image analysis techniques for the detection and classification of the early onset of plant disease and stress. Plant Methods, 2017. 13(1): p. 80.

12. Mahlein, A.K., et al., Hyperspectral Sensors and Imaging Technologies in Phytopathology: State of the Art. Annu Rev Phytopathol, 2018. 56: p. 535–558.

13. Terentev, A., et al., Current State of Hyperspectral Remote Sensing for Early Plant Disease Detection: A Review. Sensors, 2022. 22(3): p. 757.

14. Ram, B.G., et al., A systematic review of hyperspectral imaging in precision agriculture: Analysis of its current state and future prospects. Computers and Electronics in Agriculture, 2024. 222: p. 109037.

15. Zhu, H., et al., Hyperspectral Imaging for Presymptomatic Detection of Tobacco Disease with Successive Projections Algorithm and Machine-learning Classifiers. Scientific Reports, 2017. 7(1): p. 4125.

16. Zhang, X., B.A. Vinatzer, and S. Li, Hyperspectral imaging analysis for early detection of tomato bacterial leaf spot disease. Scientific Reports, 2024. 14(1): p. 27666.

17. Paulus, S. and A.-K. Mahlein, Technical workflows for hyperspectral plant image assessment and processing on the greenhouse and laboratory scale. GigaScience, 2020. 9(8).

18. Nansen, C., H. Lee, and M.B. Mesgaran, Experimental design issues associated with classifications of hyperspectral sensing data. Precision Agriculture, 2025. 26(5): p. 80.

19. Chen, Y.-N., et al., Feature Line Embedding Based on Support Vector Machine for Hyperspectral Image Classification. Remote Sensing, 2021. 13(1): p. 130.

20. Roy, S.K., et al., HybridSN: Exploring 3-D–2-D CNN Feature Hierarchy for Hyperspectral Image Classification. IEEE Geoscience and Remote Sensing Letters, 2019. 17: p. 277–281.

21. Salazar-Vazquez, J. and A. Mendez-Vazquez, A plug-and-play Hyperspectral Imaging Sensor using low-cost equipment. HardwareX, 2020. 7: p. e00087.

22. Koornneef, M. and D. Meinke, The development of Arabidopsis as a model plant. Plant J, 2010. 61(6): p. 909–21.

23. Borhan, M.H., et al., WRR4, a broad-spectrum TIR-NB-LRR gene from Arabidopsis thaliana that confers white rust resistance in transgenic oilseed Brassica crops. Mol Plant Pathol, 2010. 11(2): p. 283–91.

24. Vaswani, A., et al. Attention is All you Need. in Neural Information Processing Systems. 2017.

25. Parkes, T., From Recognition to Susceptibility: Functional characterization of Plant-specific LIM-domain containing proteins in plant-microbe interactions, in Department of Life Sciences. 2020, University of Bath: Bath, UK. p. 261.

26. Furzer, O.J., et al., An Improved Assembly of the Albugo candida Ac2V Genome Reveals the Expansion of the “CCG” Class of Effectors. Mol Plant Microbe Interact, 2022. 35(1): p. 39–48.

27. He, K., et al., Deep Residual Learning for Image Recognition. 2016. 770–778.

28. Deng, J., et al., ImageNet: a Large-Scale Hierarchical Image Database. 2009. 248–255.

29. Hong, D., et al., SpectralFormer: Rethinking Hyperspectral Image Classification With Transformers. IEEE Transactions on Geoscience and Remote Sensing, 2021. 60: p. 1–15.

30. Loshchilov, I. and F. Hutter. Decoupled Weight Decay Regularization. in International Conference on Learning Representations. 2017.

31. Benjamini, Y. and Y. Hochberg, Controlling the False Discovery Rate: A Practical and Powerful Approach to Multiple Testing. Journal of the Royal Statistical Society: Series B (Methodological), 1995. 57(1): p. 289–300.

